# The role of actin dynamics in vesicle formation during clathrin mediated endocytosis

**DOI:** 10.1101/2025.08.18.670928

**Authors:** Jie Yuan, Yen T. B. Tran, Tomasz J. Nawara, Alexa L. Mattheyses

## Abstract

Clathrin-mediated endocytosis (CME) is an important internalization route for macromolecules, lipids, and membrane receptors in eukaryotic cells. During CME, the plasma membrane invaginates and pinches off forming a clathrin coated vesicle. We previously identified heterogeneity in this process with clathrin coated vesicles forming though multiple routes including simultaneous clathrin accumulation and membrane invagination (constant curvature; CCM) as well as membrane bending after accumulation of flat clathrin (flat to curved; FTC). The architectural dynamics of vesicle formation could be influenced by osmotic or confining pressure, membrane stiffness, fluid force, or cytoskeletal arrangement. Whether these biophysical factors regulate the heterogeneity of vesicle formation dynamics is not well understood. To address this, we investigated the interconnected roles of actin and membrane tension in CME using simultaneous two-wavelength axial ratiometry (STAR) microscopy with nanometer-scale axial resolution. First, we treated Cos-7 cells with latrunculin A (LatA) to inhibit actin polymerization and found the total number of clathrin coated vesicles increased significantly, short-lifetime curved events especially. The proportion of vesicles formed following the FTC model was reduced, the membrane curved sooner after clathrin recruitment, and vesicles were less stable in the x-y plane compared to control. Next, we disrupted actin branching by inhibiting Arp2/3 with CK-869. We found an increased delay between membrane invagination and clathrin recruitment, reduced number of curved events, increased vesicle stability and an increase in the FTC model compared to control. As loss of actin filaments also reduces membrane tension, we treated Cos7 with high osmolality to decrease membrane tension and observed similar result with LatA treated group except vesicle stability stayed unchanged. This suggested the increased curved events in LatA groups may result from reduced membrane tension. We conclude actin polymerization promotes FTC while actin branching promotes vesicle formation though the CCM.

## Introduction

Clathrin mediated endocytosis (CME) is a significant internalization pathway in eukaryotic cells key to many processes, including modulating cell signaling, nutrient uptake, excitability, recycling of membrane components, and drug delivery. CME mediates the internalization of more than 20 different receptors including epidermal growth factor receptor (EGFR) and the transferrin receptor (TfR) (1). CME can be initiated by cargo binding to a plasma membrane receptor. The coat protein clathrin is recruited to the cytosolic side of the plasma membrane by an adaptor such as the AP2 complex (2). As vesicle formation continues, the clathrin coated structure (CCS) adopts an ‘Ω’ shape and move towards cytosol. Finally, the GTPase dynamin mediates scission of the CCS and releases it as a clathrin coated vesicle (CCV) into the cytosol (3). The clathrin coat is removed in a process mediated by HSP80. Many vesicles are destined for the endosome where a sorting mechanism will determine whether the cargo will be digested or returned to the plasma membrane.

Clathrin forms a triskelion structure composed of three clathrin heavy chains (CHCs) and three light chains (CLCs). When the triskelia interact, they form a polyhedral lattice that surrounds the vesicle. The CHC N-terminal domains are rich in super helix structures, which form distal bendable legs that can bend and stretch (4,5). This flexibility of the CHC allows the clathrin triskelion to form a meshwork on flat membrane (mainly hexagonal) and curved membrane (mix of hexagonal and pentagonal) (6). The distinction between flat and curved clathrin along with the observation of different clathrin architectures on the plasma membrane has led to different hypothesis of CCV formation.

Two models, constant curvature model (CCM) and flat-to-curved model (FTC, also called constant area model) are two leading models to describe the invagination of membrane during CME (7,8). In the CCM model, recruitment of clathrin happens along with curving of membrane, and the radius of curving membrane was thought to be consistent over the time. In the constant area model, clathrin accumulates as flat clathrin lattice (FCL) at the inner side of membrane until certain stage. Then the whole flat clathrin structure will bend with the membrane into a spherical dome and continue to form vesicle. While these models provide a useful framework, the formation of CCVs is more diverse and flexible. Initiation of a CCV can range from one clathrin triskelion to a FCL of varying sizes (9). Following initiation, vesicle formation can proceed through multiple pathways (10-12). Contributing to this complexity are the more than 50 different proteins reported to be involved in CME (2). For example, bin-amphiphysin-rvs (BAR) domain proteins can sense and induce membrane curvature and possibly influence the route of vesicle formation (13). Forces promoting membrane invagination during vesicle formation could be contributed by clathrin, adaptor proteins, and/or actin dynamics (7,13,14). The impact of cell type and physical environment on vesicle formation is not well understood.

The actin cytoskeleton provides a dynamic framework in eukaryotic cells, with roles including supporting and shaping the plasma membrane and mediating cell movement. Actin filaments distribute through the cells especially cell periphery and are essential for cell shape maintenance and cell movement. Polymerization and branching of actin generates force to push the membrane forward (15,16). In yeast, actin is required in the formation of clathrin coated vesicles (14,17). But due to the cell wall and turgor pressure in yeasts, results from yeast cannot represent mammalian cells. In mammalian cells, electron and fluorescence microscopy has confirmed that actin is located alongside clathrin coated pits but there is still no common agreement of the role actin plays in CME (18-20).

Understanding the role of actin in clathrin vesicle formation at single endocytic sites is necessary to distinguish the possible mechanism of local membrane bending. To address this, we applied Simultaneous Two-wavelength Axial Ratiometry (STAR) microscopy to investigate the role of actin polymerization and branching during CCV formation. STAR provides nano-scale axial resolution by exploiting the evanescent field generated by total internal reflection (21,22). Axial dynamics provide a key to distinguish different CME dynamics including whether individual vesicles are formed through CCM or FTC models. Additionally, it distinguishes if clathrin accumulations result in vesicle formation or represent dynamic flat assemblies (21,23). Here, we apply STAR to understand how disruption of actin dynamics and membrane tension impact clathrin vesicle formation. We conclude that while actin is not necessary for CME in Cos-7 cells, actin filaments stabilize vesicles during formation and actin branching promotes the generation of initial vesicle curvature.

## Materials and Methods

### Cell culture

Cos-7 cells stably expressing CLCa-iRFP713-EGFP (21) were cultured in Dulbecco’s modified Eagle’s medium (DMEM, Corning, 10013CV) containing L-glutamine and sodium pyruvate and supplemented with 10% fetal bovine serum (Gibco, 10438-026) and 2% 100 IU/ml penicillin-streptomycin (Gibco, 15070-063). For experiments, 25 mm #1.5H glass coverslips (Thorlabs, CG15XH1) were coated with fibronectin (F2006-2MG, Sigma) for 45 minutes and 60 thousand Cos-7 cells were seeded for imaging the next day.

### Cell treatments

For treatment of Latrunculin A (LatA, 428021-100UG, EMD Millipore Corp), Cos-7 cells were incubated in FluoroBrite DMEM (Gibco, A1896701) containing 5μM biliverdin (Cayman Chemical Company # 19257) and 10% fetal bovine serum (FBS) for 15 minutes. Then cells were cultured in FluoroBrite DMEM with 10% FBS, 5 μM biliverdin and 0.3 or 0.6 μM LatA for 15 minutes. 0.6μL DMSO (Fisher scientific, BP231-100) was added to mock group as control. For imaging, cells were changed to FluoroBrite DMEM (with 10% FBS and 0.3 or 0.6 μM LatA for treatment and 0.6 μL DMSO for mock) for imaging. Imaging lasted 5 minutes on each cell and 4 cells were imaged for each group.

For the treatment of CK869 (182516-25MG, EMD Millipore Corp), Cos-7 cells were starved in FluoroBrite DMEM for 30 minutes and then changed into FluoroBrite DMEM (with 5 μM biliverdin and 200 μM CK869) for 15 minutes. After that, cells were imaged with FluoroBrite DMEM (with hEGF (100 ng/ml; Millipore Sigma, E9644-2MG) and 200 μM CK869 for 5 minutes. 4 cells were imaged for each group.

Hyperosmotic medium was prepared by dissolving 0.41 grams of sucrose (Fisher S3-500) in FluoroBrite DMEM to achieve 440 mOsm/kg osmolarity. Hypoosmotic medium was prepared by mixing ddH2O (Fisher, BP2470-1) with FluoroBrite DMEM. 300 μL ddH2O in 700 μL FluoroBrite resulted in an osmolarity of 232 mOsm/kg and 600 μL ddH2O in 700 uL FluoroBrite resulted in an osmolarity of 132 mOsm/kg. The osmolality was measured by Vapor Pressure Osmometer. For hypotonic and hypertonic treatment, Cos-7 cells were starved in FluoroBrite DMEM for 30 minutes and then changed into FluoroBrite DMEM containing 5 μM biliverdin for 20 minutes. After that, cells were imaged with hyper/hypo osmotic FluoroBrite DMEM containing hEGF (100 ng/ml).

### Microscopy

Live-cell STAR image acquisition was performed with a Nikon Ti-2 microscope equipped with a motorized stage, stage-top incubator to maintain 37 °C and 5% CO_2_ (Tokai Hit, INUBG2SF-TIZB), 60x 1.49-NA objective, manual TIRF illuminator (Nikon, TI-LA-TIRF), 488 nm (Obis, 488-150 LS), and 647 nm (Obis, 1196627) excitation lasers, fiber coupling optics: fiber mount (Thorlabs, MBT621D), converging and directing the laser objective (Olympus, RMS10X), optical fiber (Thorlabs, P3-405BPM-FC-2), C-NSTORM QUAD 405/488/561/638 nm TIRF dichroic. Images were acquired with an Optosplit III (Cairn Research) image splitter with ET525/50 m and ET705/72 m emission filters (Chroma), and T562lpxtr-UF2 and T640lpxtr-UF2 dichroic mirrors to split the fluorescence emission onto separate regions of the ORCA-Flash 4.0 v3 scientific complementary metal-oxide-semiconductor camera (Hamamatsu). The system was coupled by a data acquisition device (NIDAQ, National Instruments, BNC-2115) and controlled using Nikon Elements software (version 5.02) and Coherent Connection software (version 3.0.0.8). Image acquisition was performed through NIS JOBS. Optosplit III was calibrated using the manufacturer protocol and the NanoGrid (Miraloma Tech, LLC, A00020).

### Data correction files

On every experimental day, two types of correction files were generated. (1) Cairn image splitter calibration—stack of 10 images of the NanoGrid with transmitted light illumination. This is required for image registration. (2) Flat field correction—To correct for the inhomogeneity of the TIRF excitation field, a stack of 10 images was acquired of fluorescein (ACROS ORGANICS, 2321-07-5) excited by the 488 nm laser and DiD (Invitrogen, D7757) excited by the 647 nm laser. Stock fluorescein was prepared in 1 M NaOH at 1 mg/ml and diluted at 5 μl/ml in NaOH on the day of imaging. DiD was resuspended according to manufacture instructions and diluted at 5 μl/ml in EtOH. The dilutions were applied on clean 25 mm #1.5H glass coverslips (Thorlabs, CG15XH1— coverslips were scratched in middle using a blade to find the focal plane that matches plasma membrane-coverslip interface) and mounted into Attofluor Cell Chambers (Invitrogen, A7816). TIRF images of both coverslips were taken separately with simultaneous 488 and 647 nm excitation to mimic live-cell imaging conditions.

### Image processing

Data were corrected and analyzed using Fiji (ImageJ, National Institutes of Health, Bethesda, MD), MATLAB version 2023b for DrSTAR (21,22), and CMEanalysis (24).

### Statistics and reproducibility

All live-cell experiments were performed with three independent sets of transfections, unless stated otherwise. Figure legends contain the n values for each data set, total number of cells analyzed, and description of statistical test used. All results are presented as mean ± SEM unless otherwise noted. Statistical calculations were performed in GraphPad Prism (Version 10.0.2). All not significant comparisons (P > 0.05) are not shown unless noted.

## Results

### Actin polymerization is not necessary for CME

Actin dynamically transitions between monomeric globular actin (G-actin) and filamentous actin (F-actin) (25). To test how actin polymerization impacts CME vesicle formation, Cos-7 cells stably expressing the ratiometric STAR probe clathrin light chain a (CLCa)-iRFP713-EGFP were treated with Latrunculin A (LatA). LatA binds to G-actin monomers thereby preventing the polymerization of actin filaments (26). Therefore, as actin filaments dissemble due to their native dynamics they cannot re-assemble because the G-actin is sequestered, and F-actin is lost. The disruption of actin filaments and changes to cell morphology were confirmed with phalloidin staining of F-actin and differential interference contrast (DIC) microscopy (Fig. 1A). Treatment with 0.3 or 0.6 μM LatA resulted in a similar reduction of cell attachment quantified by measuring the cell adhesion area from live cell imaging (Fig. 1B). Both concentrations were utilized in all experiments to ensure maxim disruption of actin. Clathrin puncta were observed by TIRF in control and LatA groups with co-localization of iRFP and EGFP as expected (Fig. 1C). Kymographs revealed dynamic puncta in all groups CME events, indicating actin polymerization was not necessary for CME in Cos7 cells (Fig. 1D). The live-cell imaging of iRFP and EGFP was processed by DrSTAR to generate the Δz channel which reports the curvature changes of each clathrin puncta over time (Fig. 1C and 1D). There was a range of Δz from EGFP and iRFP puncta indicating both endocytic events and flat clathrin lattices (Fig. 1C and 1D). Diffraction limited clathrin puncta that appeared and disappeared during the 5-minute experimental time frame were tracked using CMEAnalysis. The tracked puncta were separated into clathrin-coated vesicles (CCVs) and flat clathrin lattices (FCLs) depending on their dynamic Δz curvature determined from STAR (21). The shape of the clathrin structure is obtained from Δz, which is calculated as the difference between the axial position of the clathrin relative to the surrounding plasma membrane. Puncta with positive Δz, indicating an “omega” shape, were classified as CCVs and those with no change in Δz over time were classified as FCLs.

**Figure 1.**
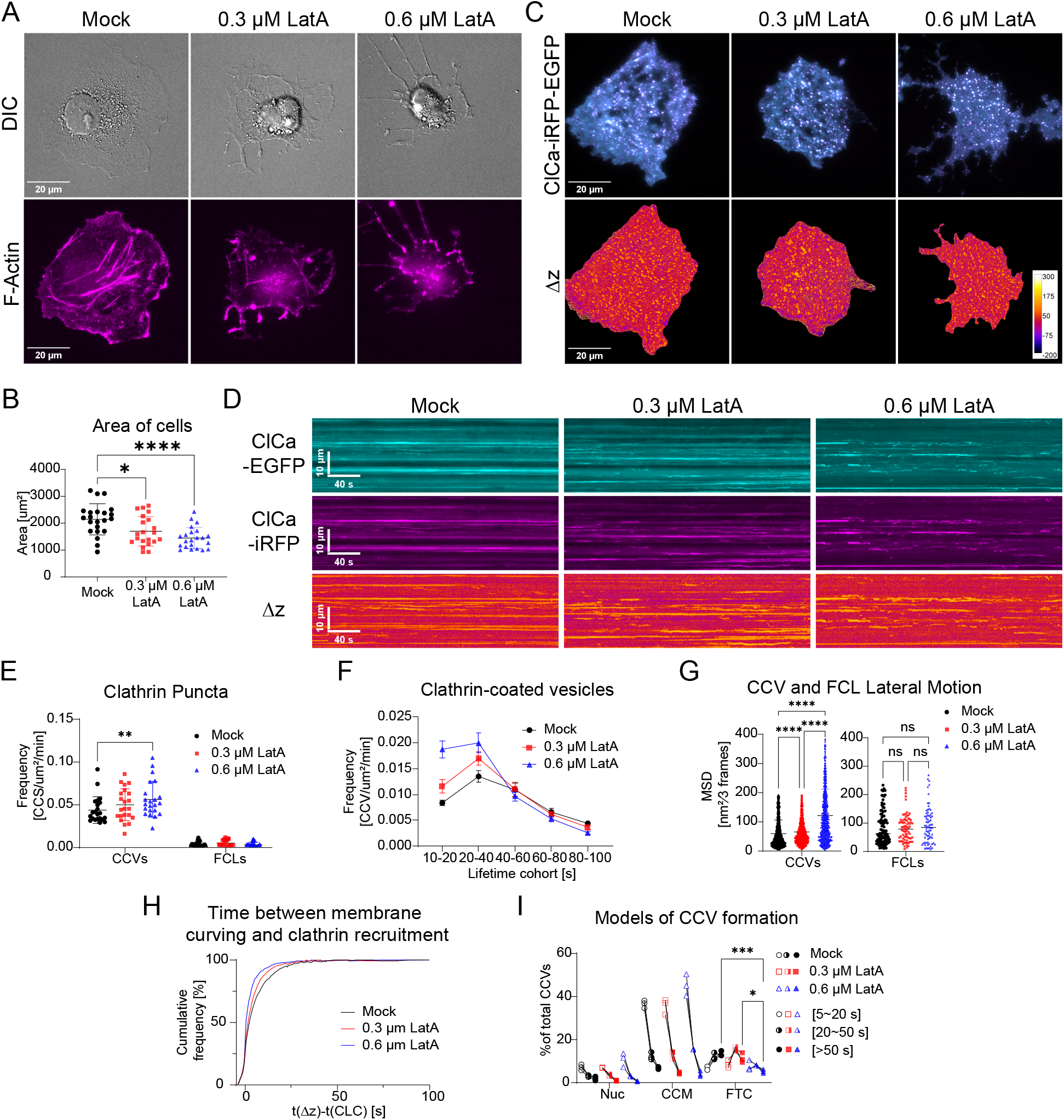
Disruption of actin filaments by LatA increased CME dynamics. (A) Representative images of Cos-7 cells stably expressing the STAR probe CLCa-iRFP713-EGFP treated with DMSO (mock), 0.3 μM or 0.6 μM LatA and stained with Phalloidin 565. (B) Cell area measured from live cell imaging of the STAR probe. 22 cells per group. mean ± SD (C) Representative images of EGFP and iRFP (overlay) and Δz from STAR microscopy. (D) Representative kymograph of EGFP, iRFP and Δz channels (E) Total density of curved clathrin vesicles (CCVs) and flat clathrin lattices (FCLs) per μm^2^, per minute. Each data point represents a cell, with 22 cells per group from 3 independent experiments; mean ± SD. Data was normally distributed by Brown-Forsythe and quantified by Welch ANOVA test. (F) Histogram of lifetime distribution of CCVs per μm^2^, per minute; mean ± SEM. (22 cells per group). (G) Motion analysis of CCVs and FCLs with a gap of three frames (0.9 seconds) by mean square displacement (MSD) with Kruskal-Wallis test. (Each data point represents a CCV or FCL. Analyzed number of CCVs: Mock = 2027, 0.3 μM LatA = 2242, and 0.6 μM LatA = 1588. FCLs: Mock = 99, 0.3 μM LatA = 79, and 0.6 μM LatA = 72. The same CCVs were analyzed in Fig. 1H,I.) (H) Cumulative histogram of the time between the initiation of membrane curvature and recruitment of clathrin. (I) Summary of CCVs across three models: Nuc = Nucleation (t(Δz) -t(CLC)< −1s), CCM = Constant Curvature Model (1 s ≤ t(Δz) -t(CLC) ≤ 4 s), FTC = Flat-to-curved transition (t(Δz) -t(CLC) > 4 s). CCVs were also classified by lifetime: 5-20 s = short-lived, 20–50 s = intermediate, and >50 s = long-lived. (* p ≤ 0.05; ** p ≤ 0.01; **** p ≤ 0.0001)

In cells treated with 0.6 μM LatA we observed an increase in the frequency of CCVs while the FCLs remained constant (Fig. 1E). Individual CCVs were next divided into cohorts based on their lifetime. This revealed the increase in the number of CCVs originated from vesicles with lifetimes between 10-40 sec, while the frequence of longer-lived (> 40 sec) CCVs remained unchanged (Fig. 1F). This suggests LatA promoted faster CCV formation, possibly leading to more CCVs. Next, we applied the mean square displacement (MSD) analysis for 2-dimensional trajectory tracking of CCVs in the x-y plane parallel to the plasma membrane. CCVs fluctuated more in the LatA group while the motion of FCLs were not altered significantly (Fig. 1G). This indicates actin filaments may have a role laterally stabilizing CCVs during CME.

Finally, we wanted to know if disruption of actin polymerization influenced the model of vesicle formation. For each CCV, we determined the delay between the initiation of vesicle formation and clathrin assembly. The time between vesicle formation and clathrin recruitment was reduced in LatA treated cells, represented by the left shift of the cumulative histogram (Fig. 1H). This suggests curvature and clathrin were more likely to initiate together when actin polymerization was disrupted. To better understand this change, we further classified the CCVs into 3 lifetime cohorts and 3 different models of vesicle formation, nucleation, CCM and FTC, based on the delay between curvature and clathrin assembly (21). If membrane curving occurred at least 1 second before clathrin detection, the CCV was classified as following the nucleation model. If membrane curving initiated at least 4 seconds after clathrin assembly, the CCV was classified as FTC model. The remaining CCVs were considered to have “no delay” between curvature and clathrin assembly (–1 to 4 sec) and were classified as following the CCM model. We found the proportion of longer lifetime CCVs formed though the FTC model decreased (Fig. 1I) while there was an increase in shorter lifetime CCM. In Cos7 cells which were serum starved treatment with a lower concertation of LatA (0.03 μM) had a smaller impact on overall cell morphology but led to similar overall results (Fig. S1). This suggests that actin may support flat clathrin, and with loss of actin there is a reduction in vesicles using the FTC model of formation. Within the same cell, we found clathrin assembly and vesicle formation occur with variable timing and actin filaments promote the flat to curved transition model. Together, this suggests actin may serve as an anchor to hold and stabilize the CCV. When actin is lost, CCVs had less lateral stability but developed curvature faster and favored the constant curvature model.

### Actin branching promotes CCV formation

The actin network contains not only linear filaments but complex branched structures as well. To test how actin branching impacts CME vesicle formation, stable Cos-7 cells expressing CLCa-iRFP-EGFP were treated with CK669. CK869 is a small molecule which binds to Arp2/3 and disrupts the nucleation of actin branching but does not affect the overall polymerization and depolymerization of actin filaments (27). As actin filaments remained intact, the cell morphology did not obviously change, and dynamic clathrin puncta were present at the membrane (Fig. 2A-D). Interestingly, in the CK869 treated group the overall frequency of CCVs was reduced, especially for the short lifetime cohorts (Fig. 2E-F). CCVs were more stable in x-y in CK869 treated group compared to control (Fig. 2G).

**Figure 2.**
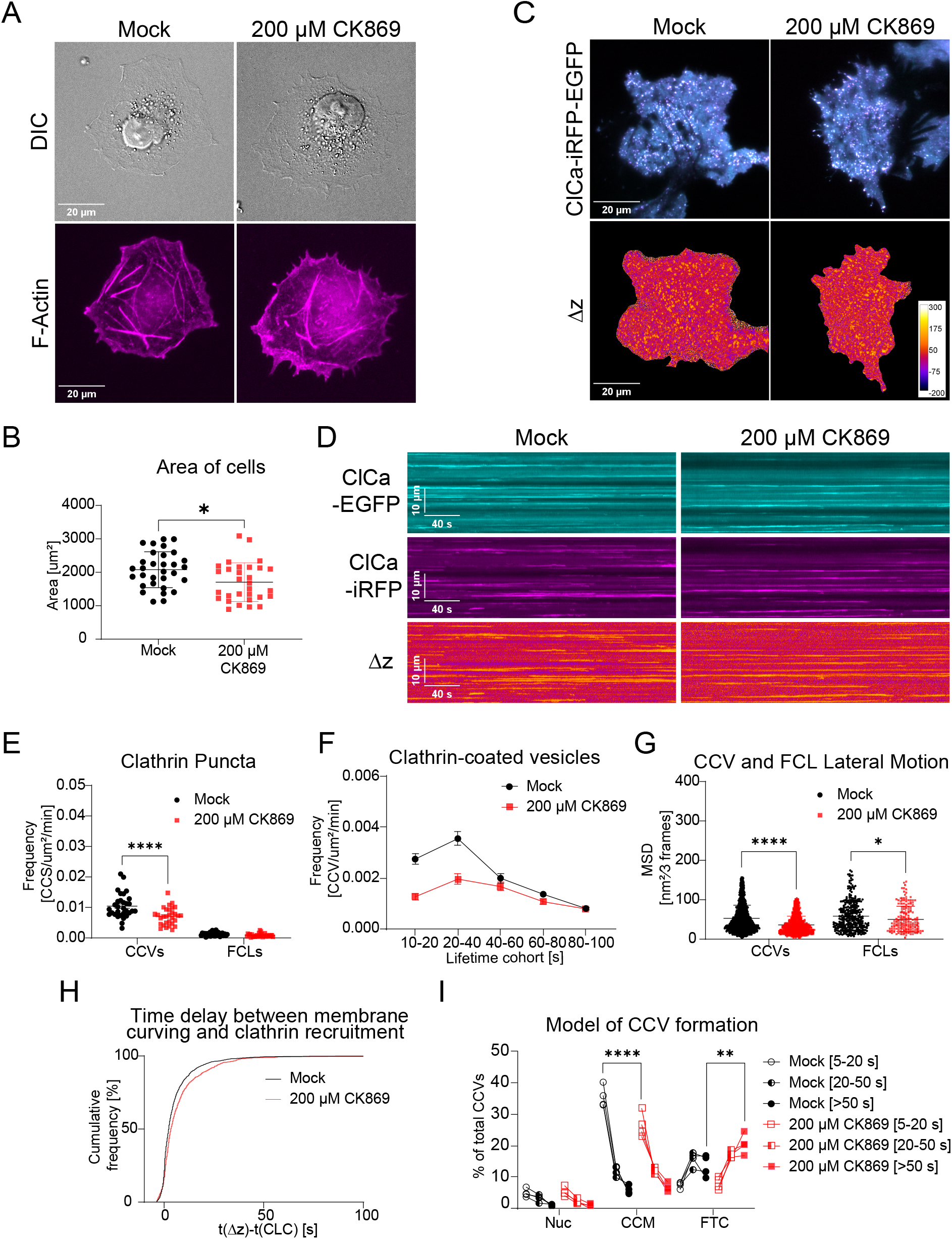
Inhibition of actin branching promotes CME dynamics. (A) Representative images of Cos7 cells stably expressing the STAR probe CLCa-iRFP713-EGFP treated with DMSO (mock) or 200μM CK869 and stained with Phalloidin 565. (B) Cell area measured from live cell imaging of the STAR probe. Mock: 31 cells, 200 μM CK869 treated: 30 cells. mean ± SD (C) Images of EGFP and iRFP overlay and Δz from STAR microscopy. Representative images from four independent replicates. (D) Kymograph of EGFP, iRFP and Δz channels. (E) Total density of CCVs and FCLs per μm^2^, per minute, mean ± SD, data was normally distributed, two-way ANOVA test was performed. (31 cells of mock, 30 cells of 200μM CK869). (F) Histogram of lifetime distribution of clathrin coated vesicles (CCVs) per μm^2^, per minute—mean ± SEM. (31 cells of mock, 30 cells of 200μM CK869) (G) Motion analysis of each condition with a gap of three frames (0.9 seconds) by mean square displacement (MSD) with Mann Whitney test. (Analyzed number of CCVs: Mock = 2299, 200μM CK869 = 1482. FCLs: Mock = 326, 200 μM CK869 = 180. The same CCVs were analyzed in Fig. 2H,I.) (H) Cumulative histogram of the time between the initiation of membrane curvature and recruitment of clathrin. (I) Summary of CCVs across three models classified as specified in Fig. 1I. (** p ≤ 0.01; **** p ≤ 0.0001)

Next, we wanted to know if disruption of actin branching influenced the model of vesicle formation. For each CCV, we determined the delay between the initiation of vesicle formation and clathrin assembly. This time was increased in CK869 treated cells, represented by the right shift of the cumulative histogram (Fig. 2H). This suggests clathrin remained in a flat configuration on the membrane for a longer time before invagination when actin branching was disrupted. The proportion of short lifetime puncta following the CCM model was reduced while the proportion of long lifetime puncta following FTC was increased (Fig. 2I). Surprisingly, LatA and CK869 had opposite effects on CME, with LatA promoting it and CK869 reducing it. This suggests that newly formed actin branches may have a different role than existing actin filament during CME. Actin branching could promote the formation of CCVs by the force generated through elongation of branches.

### Membrane tension impacts CME vesicle formation

We observed the adhesive “footprint” of the cells shrunk following LatA treatment, which could indicate reduced membrane tension (Fig. 1A). Work from Roffay et al. showed that treatment with LatA and hyper-osmotic media both resulted in a reduction of membrane tension (28). Therefore, we hypothesized that membrane tension could be a factor in our experiments, and perhaps underlies the opposing results from the LatA and CK869 treatments. To test this, we reduced membrane tension by introducing hyperosmotic media containing extra sucrose to Cos7 cells (29). There was no significant change in cell morphology and CCVs were observed in both treated and control groups (Fig. 3A-D). Overall, the frequency of CCVs was increased in the hypertonic group while the frequency of FCLs remained the same (Fig. 3E-F). The increase in frequency was in the shorter lifetime (10-40s) CCVs. We also found x-y movement of CCVs was reduced (Fig. 3G). The time between vesicle formation and clathrin recruitment was reduced and there was a decreased proportion of CCVs following the long lifetime FTC model (Fig. 3G-I). We conclude that results from 0.3 and 0.6 μM LatA represent the consequence of loss of membrane tension and disruption of F-actin.

**Figure 3.**
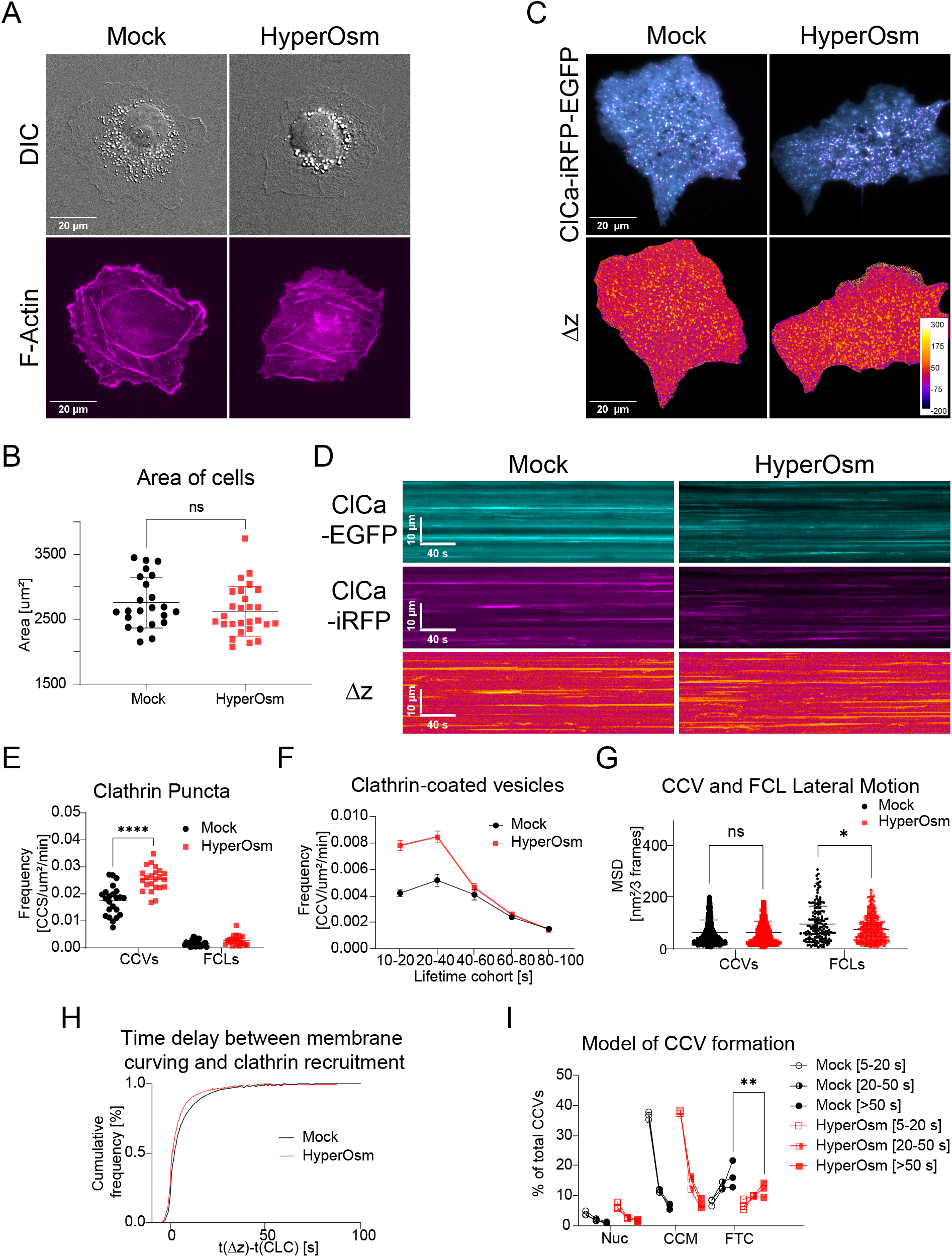
CME dynamics increase in hypertonic media. (A) Representative images of Cos7 cells stably expressing the STAR probe (CLCa-iRFP713-EGFP) treated with normal or hypertonic media (467 ± 14mOsm/kg) stained with Phalloidin 565. (B) Cell area measured from live cell imaging of the STAR probe. Mock: 23 cells, Hypertonic: 23 cells. mean ± SD (C) Overlay of EGFP and iRFP channels and Δz channel from STAR microscopy. Representative images from three replicates. (D) Kymograph of EGFP, iRFP and Δz channels. (E) Total density of CCVs and FCLs per μm^2^, per minute, mean ± SD, data was normally distributed, two-way ANOVA test is performed. (F) Histogram of lifetime distribution of clathrin coated vesicles (CCVs) per μm^2^, per minute—mean ± SEM. (23 cells of mock and 23 cells of hypertonic treated). (G) Motion analysis of each condition with the gap of three frames (in 0.9 seconds) by mean square displacement (MSD) with Mann Whitney test. (Analyzed number of CCVs: 1521 of Mock = 1521, hypertonic treated = 1350, FCLs: Mock = 172, hypertonic treated = 400. The same CCVs were analyzed in Fig. 3H,I) (H) Cumulative histogram of the time between the initiation of membrane curvature and recruitment of clathrin. (I) Summary of CCVs across three models classified as specified in Fig. 1I. (*p ≤ 0.05; ****p ≤ 0.0001)

## Discussion

Vesicle formation in CME can proceed through multiple distinct pathways. This has been shown by polarized TRIF (11), STAR (21), cryogenic electron microscopy (cryo-EM) (30) and correlative light-electron microscopy (CLEM) (12). It has been proposed that the ability to form vesicles in different ways can accommodate different cargos, be modulated by expression of accessory proteins, and/or help cells adapt to different environments. The inherent flexibility in CME could be an important underpinning for the resilience of this process required for cell survival. However, the fast dynamics and nanoscale size of CCVs makes it challenging to distinguish between different pathways and define modulators of CME pathways. Here we demonstrate that actin organization and the resulting membrane tension influence not only the frequency of endocytic events but also the pathway of vesicle formation (Fig. 4).

**Figure 4.**
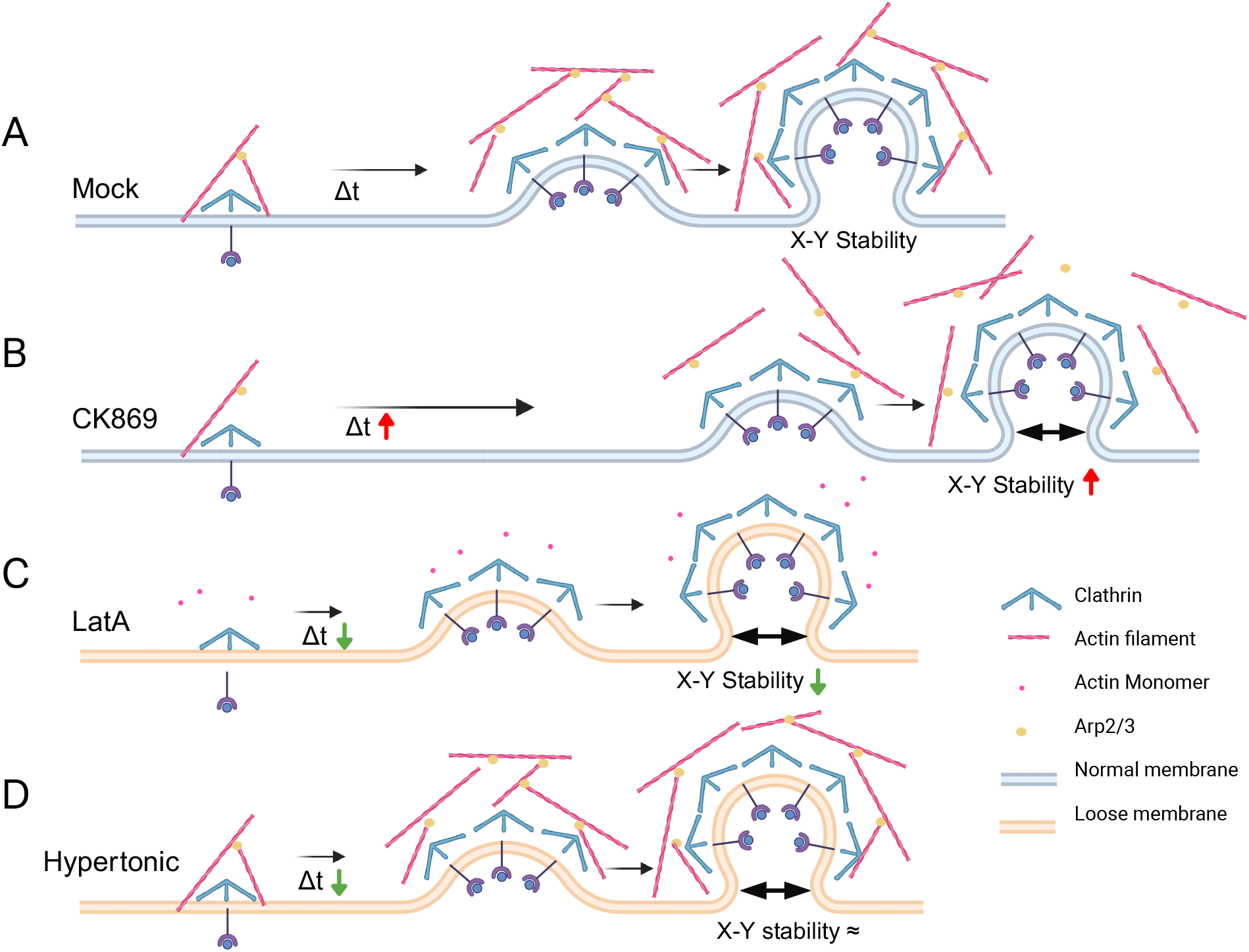
Model illustrating how actin dynamics and membrane tension impact clathrin coated vesicle formation. Illustrations depicting how actin and membrane tension can influence the dynamics of clathrin coated vesicle formation and the x-y stability of clathrin coated vesicles. (A) In mock treated cells, CCV formation can be described by the time between clathrin accumulation and formation of initial curvature and the x-y (plasma membrane plane) stability of the vesicle. (B) When actin branching was inhibited, it took longer for the membrane to form initial curvature and, the vesicle was more stable in x-y. (C) When the overall actin network was disrupted, membrane tension was reduced, the initiation of curvature happened more quickly, there was reduced x-y stability. (D) When membrane tension was reduced the initiation of curvature also happened more quickly, but the x-y stability was not affected.

One facet of CME that biophysical factors can influence the overall frequency of endocytic events. When actin polymerization was inhibited and overall actin filaments lost, there was a possibly counter-intuitive increase in the number of endocytic events. This suggests that actin polymerization is not required for successful CME in Cos7 cells, and may inhibit it. A similar increase was seen in cells treated with hyperosmotic media, suggesting the increase may be at least partially due to increased flexibility and reduced membrane tension. Interestingly, when only actin branching was inhibited, the frequency of endocytic events decreased, suggesting actin branching promotes vesicle formation. The changes in vesicle formation were primarily observed in the “shorter” lifetime events.

When actin branching was suppressed using CK869, the frequency of CCVs was reduced. The percentage of CCVs following the FTC pathway increased, suggesting forces from actin branching are not involved in this transition. Interestingly, this result also indicates that actin branching promotes early curvature form. By comparing results from hypertonic, LatA and CK869 treatment, we interpret that actin branching promoted the formation of CCVs possibly to overcome membrane tension. The ability of actin branching to promote CCV formation may come from the force generated from elongation of new branches.

Actin and membrane tension both played a role in vesicle formation. When a majority of actin structures were disrupted by LatA, accompanying the change of CME dynamics was a shrinking of the area of the cell adhered to the coverslip. The smaller footprint indicated a less tense membrane in LatA treated cells, as would be expected (28). As CME involves dramatic conformational changes of the plasma membrane, the results from LatA treatment could be affected by membrane tension. This is supported by the similarities between LatA and hypertonic treatments, which both led to an increase in the frequency of short lifetime CCVs, increased delay between recruitment of clathrin and plasma membrane curving, and a decrease in vesicles following the FTC pathway. This suggests the transition from flat to curved can be opposed by a membrane force, and by reducing this force the ability for fewer clathrin molecules to bend the membrane as they are recruited is increased. Finally, LatA but not hypertonic treatment decreased the x-y stability of CCVs during formation. This suggests actin plays a role in stabilizing CCVs during formation, either through a direct interaction or indirectly by the linkages between actin and other membrane proteins.

In all cases the frequency of FCLs remained unchanged. If FCLs represented failed endocytosis we would expect less of them in conditions that favor endocytosis and more of them in conditions that inhibit endocytosis. The data presented here suggests that FCLs do not represent “failed” endocytosis, as their frequency was not changed even with increased or decreased membrane tension. FCLs as defined here represent dynamic clathrin which appears and disappears within the imaging experiment but does not have a significant change in z position relative to the surrounding membrane. Importantly, it does not represent more static clathrin structures with longer lifetimes. These dynamic FCLs may represent an alternate role for clathrin at the plasma membrane that is not influenced by actin or membrane tension. Our data suggests they do not represent frustrated endocytosis.

The opposing results of LatA and CK869 treatment suggests that when most actin filaments were disrupted by LatA the effect from the loss of membrane tension overwhelmed the effects from loss of actin structures. We hypothesize that in physical conditions that are less favorable for vesicle formation, such as increased membrane tension, actin can act to overcome this and promote membrane bending. The local membrane environment can also be heterogeneous which can lead to higher viscosity in some membrane regions (31). This type of local heterogeneity where the membrane could be more easily deformed may contribute to endocytic “hot spots” (32). As the experiments presented here were performed in the Cos7 fibroblast cell line, they may not represent all cell types. Different cell types reside in distinguishing environments and carry out different roles. Organization of actin varies across different cell lines and cell under different conditions (33-35). Actin filaments in normal fibroblast cells are more aligned and bundled compared with neoplastically transformed cells (33). Previously we found that cultivating human umbilical cord endothelial cells under flow led to large scale changes in actin organization, enhanced endocytosis, and increased the proportion of vesicles undergoing FTC (36). Therefore there will be a difference in compose of membrane lipid, proteins and carbohydrate leading to different fluidity and stiffness of membrane (37). We conclude that actin filaments anchor and stabilize CCVs and that actin branching can promote the formation of CCVs but overall actin is not necessary for successful vesicle formation. This work contributes to the current discussion of the forces needed to overcome the unfavorable energy to form CCVs.

## Author Contributions

Conceptualization: J.Y., T.N., A.M.; Methodology: J.Y., Y.T., T.N., A.M.; Validation: J.Y., Y.T., T.N.; Formal Analysis: J.Y., Y.T.; Investigation: J.Y., Y.T., T.N.; Resources: A.M.; Writing-Original Draft: J.Y.; Writing-Review and Editing: J.Y., Y.T., T.N., A.M.; Visualization: J.Y., Y.T.; Supervision: A.M.; Funding Acquisition: A.M., T.N., and J.Y.

