## Supplemental Figures for "The role of actin dynamics in vesicle formation during clathrin mediated endocytosis"

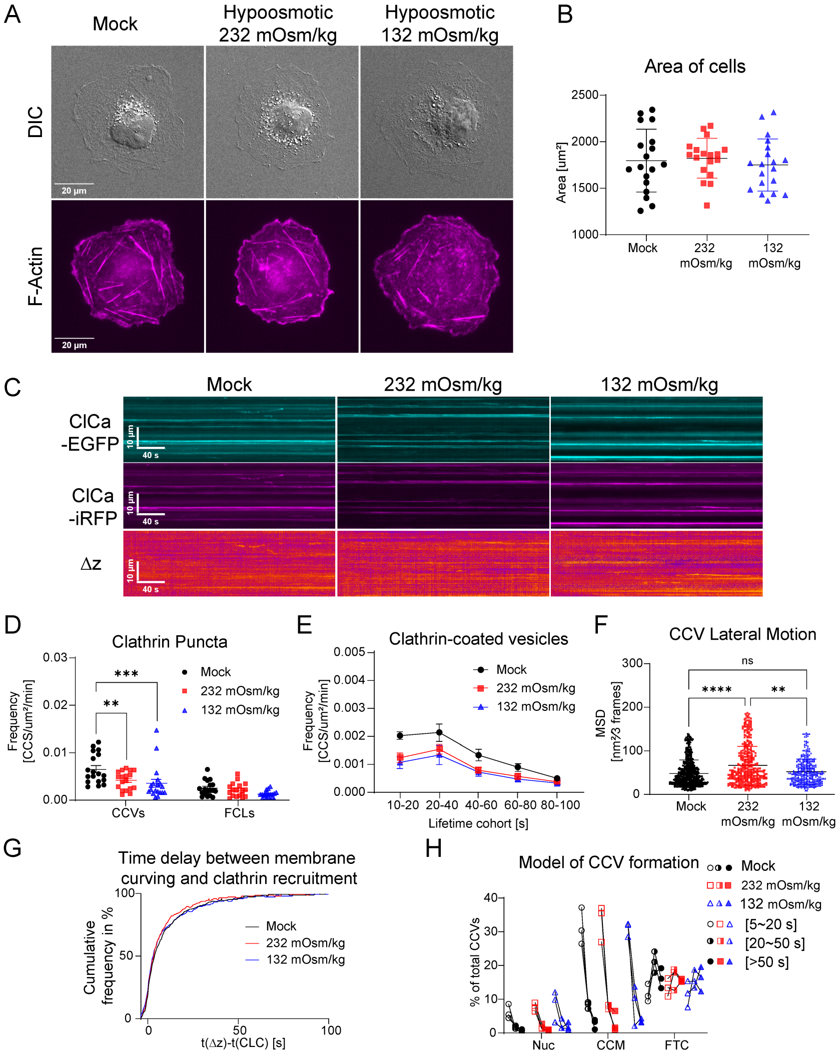


**Figure S1 CME dynamics decrease in hypotonic media** (A) Stable Cos7 cells expressing the STAR probe CLCa-iRFP713-EGFP treated with normal or hypotonic media (with an osmotic pressure of 232 mOsm/kg or 132 mOsm/kg) stained with Phalloidin 565. (B) Cell area measured from live cell imaging of the STAR probe. 18 cells for mock, 18 cells for 232 mOsm/kg, and 17 cells for 132 mOsm/kg. mean ± SD (C) Kymographs of EGFP, iRFP and Δz channels for control (Mock), Osmotic pressure 232 mOsm/kg, and 132 mOsm/kg. (D) Frequency of CCV per $\mu m^2$, per minute, mean ± SD, data was normally distributed, Two-way ANOVA. (E) Histogram of lifetime distribution of clathrin coated vesicle (CCV) and flat clathrin lattice (FCL) per $\mu m^2$, per minute—mean ± SEM. (Data in C and D, 18 cells for mock, 18 cells for 232 mOsm/kg, and 17 cells for 132 mOsm/kg from three independent repeats. (F) Motion analysis of CCVs from each condition with a gap of three frames (0.9 secs) by mean square displacement (MSD) with Mann Whitney test. (Analyzed number of CCVs: 528 of Mock, 407 of 232 mOsm/kg and 314 of 132 mOsm/kg). (G) Cumulative histogram of the time between the initiation of membrane curvature and recruitment of clathrin. (H) Summary of CCVs across three models classified as specified in Fig. 1I. (*$p \leq 0.05$; **$p \leq 0.01$; ****$p \leq 0.0001$)

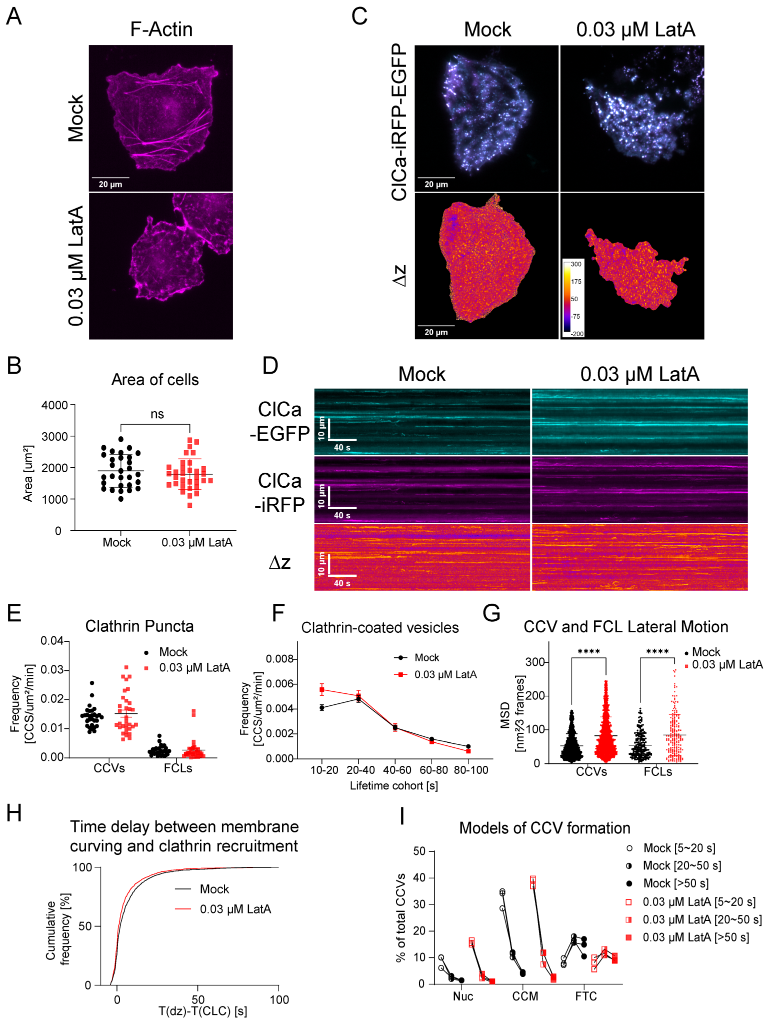


**Figure S2 Change of CME dynamics after inhibition of actin in starved Cos7 cells** (A) Stable Cos-7 cells expressing the STAR probe CLCa-iRFP713-EGFP treated with DMSO, 0.03 μM LatA and stained with Phalloidin 565. (B) Cell area measured from live cell imaging of the STAR probe. 29 cells for mock and 32 cells for 0.03 μM LatA treated (C) Registered overlay of EGFP and iRFP channels and Δz channel from STAR microscopy. Representative images from three replicates. (D) Kymograph of EGFP, iRFP and Δz channels for control (Mock) and 0.03 μM LatA. (E) Frequency of CCVs per μm$^2$, per minute (29 cells for mock and 32 cells for 0.03 μM LatA). mean ± SD per cell. Data was normally distributed Brown-Forsythe and Welch ANOVA test is performed. F) Histogram of lifetime distribution of clathrin coated vesicle (CCV) and flat clathrin lattice (FCL) per μm$^2$, per minute—mean ± SEM. (29 cells for mock and 32 cells for 0.03 μM LatA). (G) Motion analysis of CCV in each condition with the gap of three frames (0.9 secs) by mean square displacement (MSD) with Kruskal-Wallis test. (Analyzed number of CCVs: Mock = 2049, 0.03 μM LatA = 2062, FCLs: Mock = 217, 0.03 μM LatA = 196) (H) Cumulative histogram of the time between the initiation of membrane curvature and recruitment of clathrin. (I) Summary of CCVs across three models classified as specified in Fig. 1I. (****p ≤ 0.0001)
